# Optimizing resource allocation in *Miscanthus* breeding with sparse testing designs for genomic prediction

**DOI:** 10.64898/2026.03.18.712722

**Authors:** Shatabdi Proma, Nelson Lubanga, Erik Sacks, Andrew D.B. Leakey, Hua Zhao, Bimal Kumar Ghimire, Alexander E. Lipka, Joyce N. Njuguna, Chang Yeon Yu, Eun Soo Seong, Ji Hye Yoo, Hironori Nagano, Kossonou G. Anzoua, Toshihiko Yamada, Pavel Chebukin, Xiaoli Jin, Lindsay V. Clark, Karen Koefoed Petersen, Junhua Peng, Andrey Sabitov, Elena Dzyubenko, Nicolay Dzyubenko, Katarzyna Glowacka, Moyses Nascimento, Ana Carolina Campana Nascimento, Maria S. Dwiyanti, Larisa Bagment, Ansari Shaik, Julian Garcia-Abadillo, Diego Jarquin

## Abstract

Phenotyping high-biomass perennial crops is laborious and the rate of genetic gain in perennial crop breeding programs is typically low. So, it is especially important to identify methods that produce efficiency gains in the breeding process. *Miscanthus* is a C4 perennial grass with favorable characteristics for producing biomass as a feedstock for biofuels and diverse biobased products. Increasing biomass yield will increase profitability and environmental benefits, so is a key target for *Miscanthus* breeding. In addition, the identification of well-adapted genotypes across a wide range of environmental conditions requires the establishment of multi-environment trials (METs).

Sparse testing is a genomic prediction-based strategy that reduces the phenotyping costs in METs by selecting a subset of genotypes to evaluate in a subset of environments and then predicts the performance of the unobserved genotype-environment combinations. A *Miscanthus sacchariflorus* (MSA) population comprising 336 genotypes observed across three environments was analyzed. Three prediction models considering main effects (environments, genotypes, genomic) and interaction effects (genotype-by-environment; G×E interaction) were implemented for forecasting dry biomass yield (YDY), total culm (TCM), average internode length (AIL), and culm node number (CNN). Multiple calibration sets based on different compositions and sizes were considered to evaluate performance in terms of the predictive ability (PA) and the mean square error (MSE) for a fixed testing set size. The training set size ranged from 52 to 112 to predict a fixed set of 224 unobserved genotypes across all three environments.

The results showed that the model accounting for G×E interaction presented the highest PA and the lowest MSE for CNN (PA: ∼0.77, MSE: ∼0.5) and YDY (PA: ∼0.70, MSE: ∼1.3) while for TCM and AIL these ranged from ∼0.28 to 0.41 and ∼1.3 to 4.3, respectively. Overall, varying training sets and allocation strategies did not affect PA and MSE, with 52 non-overlapping and 0 overlapping genotypes per environment as the optimal cost-effective allocation framework. This suggests that implementing sparse testing designs could significantly reduce phenotyping costs by fivefold, without compromising PA in breeding programs for perennial crops such as *Miscanthus*.

## Introduction

Biomass crops offer a source of feedstock that can be used to help meet demand for clean fuels and diverse biobased products, while also improving soil health and water quality (Metzger and Hüttermann, 2009; Ladanai and Vinterbäck, 2009, Saha et al., 2013; Kreig et al., 2019; Sutton et al., 2025). *Miscanthus* is a perennial species that holds the potential as a biomass grass for a wide range of applications (Chung & Kim, 2012). It can be used to produce lignocellulosic ethanol, generation of electricity and heat through gasification or direct combustion, and for the manufacture of paper, construction materials, biodegradable plastics, animal bedding, mulch, and livestock feed (Clifton-Brown et al., 2001; Clifton-Brown and Lewandowski, 2002).

Currently, commercial *Miscanthus* biomass production in Europe and North America relies predominantly on a single sterile triploid clone of *Miscanthus* × giganteus (M.× g), an interspecific cross between *Miscanthus sacchariflorus* (MSA) and *Miscanthus sinensis* (MSI) (Hodkinson and Renvoize, 2001). However, the standard clone M.× g has insufficient winter hardiness in northern Europe and the northern US Midwest (Clifton-Brown & Lewandowski, 2000; Dong et al., 2019). Moreover, cultivating a single clone on a large-scale basis faces risks associated with pests and diseases (Njuguna et al., 2023). Therefore, new high-yielding and climate-resilient *Miscanthus* varieties must be developed to withstand the ever-changing climate.

There are a set of challenges common to breeding highly productive, perennial crops that must be addressed to fully exploit their beneficial characteristics, regardless of whether they are grasses or woody species. *Miscanthus* is a valuable model system in which to address these issues. *Miscanthus* breeding is challenging relative to annual crops because the crop has to establish for 2-3 years before high-quality yield data can be gathered (Chung & Kim, 2012). The large size of the crop makes phenotyping challenging, although high-throughput remote-sensing based techniques are being developed to alleviate this bottleneck (Varela et al., 2022; Varela et al., 2025).

The initial cost of establishing *Miscanthus* is also significant because the high yielding, commercial varieties are vegetatively propagated via rhizome (Lewandowski et al., 2003; Anzoua et al., 2011). Multi-environmental trials (METs) are important in *Miscanthus* breeding programs to identify superior genotypes that are both locally adapted and perform well across environments (Njuguna et al., 2023). However, field evaluation of the *Miscanthus* genotypes across multiple environments further increases costs and labor. To optimize both phenotyping cost and the accurate selection of improved cultivars, efficient approaches that can improve the efficiency in *Miscanthus* breeding programs need to be implemented (Aoun et al., 2021).

Genomic selection (GS), i.e. employing genomic prediction (GP) models in practical selection, has emerged as a powerful tool in plant breeding owing to the advancement of high-throughput sequencing and the availability of genome-wide single-nucleotide polymorphisms (SNPs). The concept of GS was introduced by Meuwissen et al., (2001) to overcome the challenges posed by having a larger number of predictors (*p)* compared to the records available (*n*) for model fitting. However, Bernardo (1994) was the first who proposed to use genomic data in the form of Restriction Fragment Length Polymorphism (RLFP) for predicting the performance of unobserved maize crosses. GS uses a GP model utilizing whole genome marker data to predict the performance of the unobserved individuals via their genomic profiles only (McGaugh et al., 2021; Persa et al., 2021).

In GP, phenotypic and genotypic data are applied to calibrate and optimize various models (Jarquin et al., 2021). Usually, before embarking on using GS in real scenarios, the levels of PA that can be reached with the available data are estimated by considering cross-validation studies mimicking realistic prediction scenarios (Cheng et al., 2021; McGaugh et al., 2021). For this, all the available data is split into two non-overlapping partitions, training and testing sets and the former is used to predict the latter partition. The training set is composed of individuals who have both phenotypic and genomic information, while the testing set contains untested individuals with only genomic information (Isidro et al., 2015). Then, the potential of untested genotypes in the testing set is evaluated.

Since the development of improved cultivars requires the establishment of METs to evaluate stability and local adaptation, it is necessary to consider models that accurately capture differences among genotypes in their response to environmental stimuli (Rincent et al., 2017). In most cases, the response patterns (rankings) change from one set of environmental conditions to another for the same group of genotypes (Crossa et al., 2004). This change in the response patterns, when it occurs, is referred as the presence of the genotype-by-environment (G×E) interaction (Jarquin et al., 2021). The breeding process can be sped up by ensuring an accurate selection of the best performing genotypes in advance while skipping field testing. Thus, integrating G×E into the GP models holds the potential to speed up the breeding process by ensuring an accurate selection of genotypes while decreasing the number of individuals to be evaluated in METs.

Previous studies have shown the incorporation of G×E interactions into GP models could potentially improve predictive ability. For example, Jarquin et al., (2014) proposed a model that implicitly allows the inclusion of the interaction between each maker SNP and each environmental covariate or environment via covariance structures. Meanwhile, Lopez-Cruz et al., (2015) presented a model that explicitly allows the incorporation of all contrasts between marker SNPs and environments. The prediction accuracies of the models are assessed using cross-validation (CV) schemes where the datasets are conveniently divided into training and testing populations mimicking real prediction problems of interest to breeders. Two of the main cross-validation schemes utilized are CV1 and CV2. CV1 predicts novel genotypes that were never tested in any environment, while CV2 predicts observed genotypes in incomplete field trials.

Sparse testing (Jarquin et al., 2020) is an innovative augmentation for field evaluation to capture G×E in METs. It can increase the testing capacity of the genotypes at a fixed cost by allowing more genotypes to be tested in target environments. Alternatively, it also offers the possibility of reducing phenotyping costs for a given set of genotypes and environments of interest (Garcia-Abadillo et al., 2024). In sparse testing, genotypes are split across environments and analyzed using different GP models to estimate their specific performance. The best design is determined by evaluating the relationship between the prediction accuracies of the models and the testing capacity of the designs. In other words, the best sparse testing design quantifies the accuracy of GP models with limited testing efforts to achieve acceptable predictive ability, ensuring efficient resource allocation without sacrificing model performance.

The optimal designs in sparse testing balance the trade-off between high accuracy, reduced training set size and diverse composition. This is achieved by calibrating training sets either by using a large set of overlapped genotypes in a few environments or sets of few common genotypes across multiple environments. The accuracy of predicting unobserved genotypes in METs is influenced by *a*) the number of overlapping genotypes across environments, *b*) the number of environments where each genotype is observed, *c*) the implemented prediction model, and *d*) the average number of phenotypes per genotype (Garcia-Abadillo et al., 2024). Sparse testing has been implemented in different plant breeding programs in major crops including maize (Jarquin et al., 2020), wheat (Atanda et al., 2022), soybean (Persa et al., 2023) and sugarcane (Garcia-Abadillo et al., 2024).

The main aims of this study are to *1*) explore the potential of implementing sparse testing designs in multi-environment *Miscanthus* trials and, *2*) compare the prediction accuracies of three prediction models. The implemented metrics for model evaluation were the correlation between predicted and observed values (PA) and the mean square error (MSE). Earlier studies have focused on using traditional GP approaches in MSA and MSI germplasm panels (Dong et al., 2019; Njuguna et al., 2023). To our knowledge, the use of sparse testing designs has not yet been explored in *Miscanthus* breeding programs.

This study considered a diverse panel of MSA genotypes in different sparse testing designs similar to those proposed by Jarquin et al., (2020). Three types of designs including two extreme allocation strategies were considered *i*) non-overlapping genotypes (NO) where each genotype is uniquely observed across environments, *ii*) overlapping genotypes (O) where all genotypes are observed across all environments, *iii*) mixture of both NO and O scenarios. In addition, different reduced training set sizes were also considered along different compositions.

For all combinations between training set sizes and compositions, three prediction models were implemented: *i*) **M1**- includes the main effects of environment and genotype and it is based mainly in phenotypic information where no marker information is used, *ii*) **M2**- includes the main effects of environment, genotype, and markers, and *iii*) **M3**- includes the main effects of environment, genotype, and marker and the interaction between markers and environments.

Overall, the study shows that a GP-based sparse testing approach can be used to increase the selection accuracy and reduce the cost of evaluating *Miscanthus* genotypes in multiple environments. This was achieved using G×E model (M3), which achieved the best performance, with the highest PA and the lowest MSE. Additionally, across all non-overlapping and overlapping design compositions, the performance of the model remained constant. This indicates that whether *Miscanthus* breeders choose to evaluate similar genotypes across various locations or different genotypes across locations, they would not have to compromise the accuracy of the model.

## Materials and methods

### Phenotypic data

The original phenotypic dataset consisted of 590 MSA accessions collected in East Asia and established in four locations [Sapporo, Japan by Hokkaido University (HU); Urbana, Illinois by the University of Illinois (UI); Chuncheon, South Korea by Kangwon National University (KNU) and Zhuji, China by Zhejiang University (ZJU)] between 2016 and 2017 as described by Njuguna et al. (2023). Briefly, productivity was phenotyped by assessing above-ground dry biomass and 15 yield component traits at every location between two and three years after planting. Only for dry biomass, a fourth-year data was also collected at all locations resulting in three years of phenotypic information for this trait. The best linear unbiased estimates (BLUES) of each trait for each genotype in every location (i.e., across years) were provided along with the dataset. These values were used to implement the sparse testing designs. Further details of how the BLUES were obtained can be found in Njuguna et al. (2023).

Out of the 16 initial traits comprising yield and its component traits only four [dry biomass (YDY), culm node number (CNN), total culms (TCM), and average internode length (AIL)] were considered for analyses due to complete information of traits across environments. In sparse testing studies, at least for demonstration purposes only, a balanced dataset is initially required (a set of common genotypes observed across locations in this case). Due to a high frequency of missing values, one of the environments (UI) was discarded from the study. After discarding genotypes with missing information for at least one trait-in-environment combination, a total of 336 individuals remained in the analyses.

### Genotyping and quality control

A detailed description of the genotyping process can be found in Clark et al., (2014). Briefly, genotyping was performed using restriction site-associated DNA sequencing (RAD-seq) following the library preparation. Hence, the library samples were sequenced with Illumina Hiseq 2500 and 4000. Raw sequence reads were aligned to the MSI reference genome and SNP calling was performed using TASSEL-GBS pipeline (Bradbury et al., 2007; Mitros et al., 2020). Initially, SNPs with more than 70% call rate were removed resulting in 268,109 SNPs. Further filtering involved removing SNPs with more than 50% missing values and a minor allele frequencies (MAF) smaller than 3%, resulting in a total of 136,814 SNPs that were used in this study.

### Allocation Scenarios

In a practical plant breeding program, only a subset of genotype-in-environment combinations can be evaluated in METs due to budget limitations and seed availability. Sparse testing designs contribute to implementing strategies in allocating genotypes across environments that extrapolate the whole population (Jarquin et al., 2020; Garcia-Abadillo et al., 2024). To implement the sparse testing strategies, the objective entails creating a training population to accurately predict the performance of the unobserved genotype-in-environment combinations using a subset of these combinations observed in fields as calibration set. In general, according to their composition the calibration sets can be classified in the following types:

**A. *Overlapping (O) genotypes:*** It resembles the original concept of genomic selection of predicting the performance of unobserved or newly developed genotypes in one or several environments where other genotypes were already observed. Usually, this manner to partition the available information in training and testing sets is also known as CV1 scheme.
**B. *Non-overlapping (NO) genotypes:*** This design fits perfectly in the METs implementation where some genotypes are observed in some environments but not in others in a sparse arrangement. In principle, this manner of composing training sets is regarded as CV2 scheme. The most extreme case considers observing only once each genotype across environments resulting in the non-overlapping design.
**C. *No-overlapping/Overlapping genotypes:*** Alternative designs can be form combining sets of overlapping and non-overlapping genotypes. Hence, the original two designs can be modified into different fractions of NO/O genotypes.

In the present study, considering a total population of 336 genotypes, assessed in three environments, provides 1,008 phenotypes (336 x 3). With this data, three different types of designs were implemented, all of which had a fixed testing set size of 224 genotypes per environment (i.e. 672 (224×3) phenotypes across the three environments). Correspondingly, the training set size varied from a maximum of 112 genotypes observed per environment (i.e. 336 (112 x 3) phenotypes across the three environments) to a minimum of 52 genotypes per environment (i.e. 156 (52 x 3) phenotypes across the three environments). TRAINING set sizes were varied in step sizes of 10 between 112 and 52 (e.g. 102, 92, 82, 72, and 62).

**1. *Completely non-overlapping allocation design (NO/O = 112/0 genotypes)***: In this design, 112 non-overlapping genotypes were observed in each environment (i.e. 112 genotypes uniquely observed in one environment), with zero overlapping genotypes (i.e. 0 genotypes assessed in all three environments).
**2. *Completely overlapping allocation design (NO/O = 0/112 genotypes):*** In this design, 0 non-overlapping genotypes were observed in each environment (i.e. 0 genotypes uniquely observed in one environment), with between 52 and 112 overlapping genotypes (i.e. between 52 and 112 genotypes assessed in all three environments).
**3. *Mixed allocation design (NO/O = 102/10 varying to 2/110 genotypes:*** In these designs, data included a mixture of phenotypic information for non-overlapping genotypes that were observed in a single environment and overlapping genotypes that were observed in all three environments.

Many possible combinations of non-overlapping and overlapping genotypes could be used, given the available dataset; however, sets of 10 common genotypes were conveniently traded to study prediction patterns of the training set composition in PA. Assuming that *C* represents the common set of genotypes across the three environments, *U* is the unique set of genotypes that are different across the three environments, and *L* is the number of environments. Then, the total of genotypes-in-environment combinations in the calibration set is (*L*×*C*) + (*L*×*U*) = [*L*×(*C*+*U*)] (Garcia-Abadillo et al., 2024).

In this study, the calibration sets were constructed to include the mixing extremes of XX/0, 0/XX, and mixed type XY/XW in function of different per environment sample sizes varying as follows 112, 102, 92, 82, 72, 62, and 52. The details of the different designs can be found in Table 1 where the rows represent the training set size and columns present different combinations or levels of non-overlapping/overlapping sets. For each specific calibration size (size and composition), 10 random re-samplings of genotype selections were considered. The different training set compositions can be visualized in Appendix Figure S1.

**Table 1.**
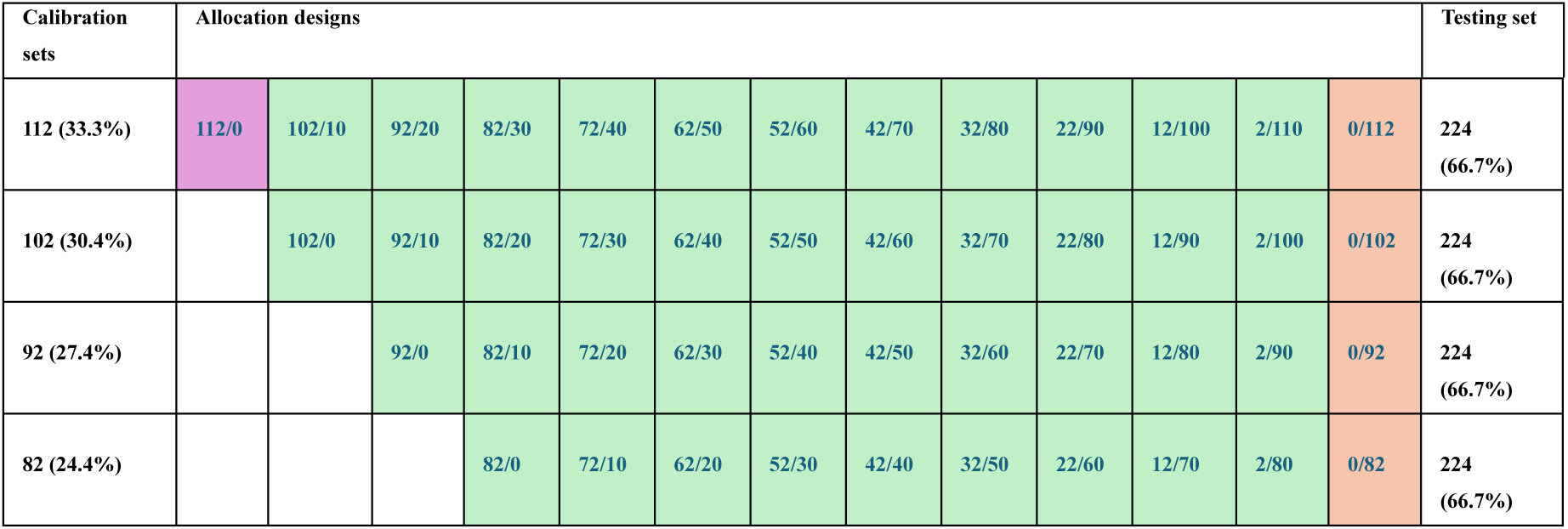
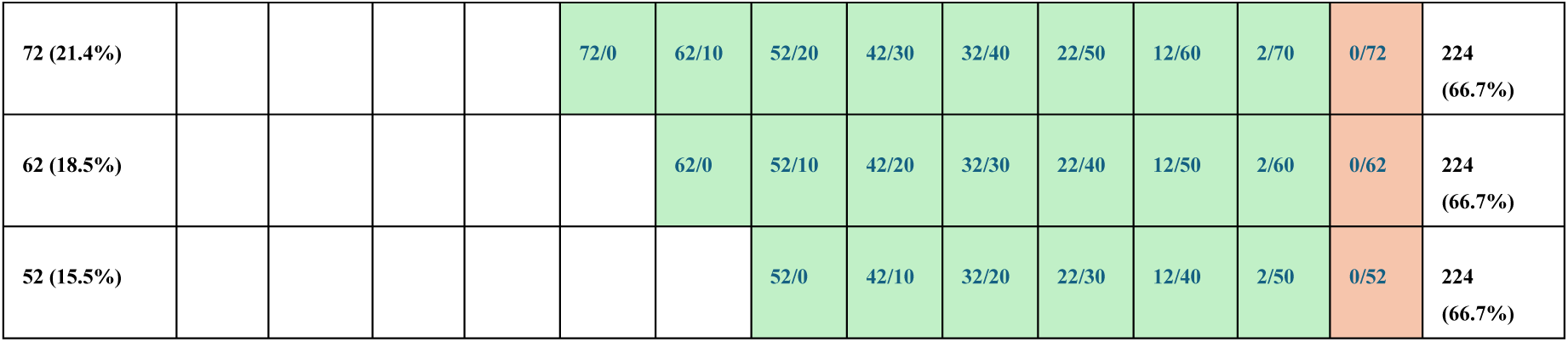
Combinations of non-overlapping/overlapping (NO/O) allocation strategies for different calibration set sizes. The column in the far left presents different sampling sizes (and percentages with respect to all the potential genotype-in-environment combinations). The column in the far right presents the fixed within-environments prediction set size (and percentage). The columns in the middle show the different allocation designs as a function of the training set size. Completely overlapping designs are shown in orange cells. The completely non-overlapping design is shown in a purple cell. Mixed allocation designs are in green cells.

| Calibration sets | Allocation designs |  |  |  |  |  |  |  |  |  |  |  |  | Testing set |
| --- | --- | --- | --- | --- | --- | --- | --- | --- | --- | --- | --- | --- | --- | --- |
| 112 (33.3%) | 112/0 | 102/10 | 92/20 | 82/30 | 72/40 | 62/50 | 52/60 | 42/70 | 32/80 | 22/90 | 12/100 | 2/110 | 0/112 | 224 (66.7%) |
| 102 (30.4%) |  | 102/0 | 92/10 | 82/20 | 72/30 | 62/40 | 52/50 | 42/60 | 32/70 | 22/80 | 12/90 | 2/100 | 0/102 | 224 (66.7%) |
| 92 (27.4%) |  |  | 92/0 | 82/10 | 72/20 | 62/30 | 52/40 | 42/50 | 32/60 | 22/70 | 12/80 | 2/90 | 0/92 | 224 (66.7%) |
| 82 (24.4%) |  |  |  | 82/0 | 72/10 | 62/20 | 52/30 | 42/40 | 32/50 | 22/60 | 12/70 | 2/80 | 0/82 | 224 (66.7%) |
| 72 (21.4%) |  |  |  |  | 72/0 | 62/10 | 52/20 | 42/30 | 32/40 | 22/50 | 12/60 | 2/70 | 0/72 | 224<br>(66.7%) |
| 62 (18.5%) |  |  |  |  |  | 62/0 | 52/10 | 42/20 | 32/30 | 22/40 | 12/50 | 2/60 | 0/62 | 224<br>(66.7%) |
| 52 (15.5%) |  |  |  |  |  |  | 52/0 | 42/10 | 32/20 | 22/30 | 12/40 | 2/50 | 0/52 | 224<br>(66.7%) |

### Prediction models

Three random effects prediction models were considered, one based on phenotypic information only and the other two involving genomic information as main and interaction effects (Jarquin et al., 2020; Garcia-Abadillo et al., 2024). **M1: E+L**, is the base model that considers the genotype and environment main effects only. These effects are assumed to follow independent and identical (*iid*) normal distributions. Hence, in this case, there is not a mean to connect calibration and testing sets when the phenotypic information is absent for genotypes across environments.

**M2: E+L+G**, extends the M1 model by including the marker information as a main effect via covariance structures for **G**. This model allows the borrowing of information between training and testing for never-observed genotypes. One expected drawback of this model is that estimated genomic values of a particular genotype across environments are identical, impeding modeling of phenotypic variation across different environments on a genotype-specific basis.

**M3: (E+L+G+GE)** is the extension of M2 with the inclusion of the genomic-by-environment (G×E) interaction term. This model allows genotype-specific responses in each environment, and these are simulated with a common genomic effect or intercept plus a gradient slope modeled by the interaction effect.

The performance of the models was evaluated for all training set composition and sizes. The metrics considered are the PA [as the Pearson correlation coefficient between observed and predicted values] and the MSE [mean of the squared of the differences between observed and predicted values]. Further details of these models are provided below.

### M1: E+L [Environment + genotype]

Consider that *y*_*ij*_ represents the adjusted phenotypic observation of the *i*^*th*^ (*i* = 1,2, …, *L*) genotype in the *j*^*th*^ (*j* = 1,2, …, *J*) environment and it is explained as the sum of the common mean *μ*, the random effect of the *j*^*th*^ environment (*E*_*j*_), the random effect of the *i*^*th*^ genotype (*L*_*i*_), plus a random error term (*ɛ*_*ij*_). Further assumptions are made regarding the relationships among the random terms *E*_*j*_, *L*_*i*_ and *ɛ*_*ij*_ considering these independent between them. In addition, for each random term their corresponding levels are considered independent and identical normal distributed (*iid*). Hence, the model can be written as follows

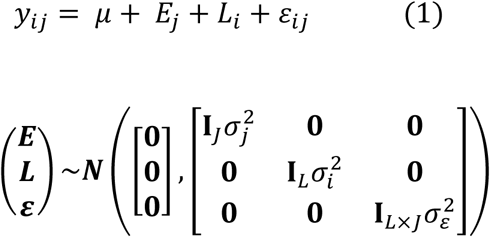

where **I**_*J*_, and **I**_*L*_ are the identity matrices according to the levels of environments and genotypes.

### M2: E+L+G [environment + genotype + genomic (markers)]

M2 is an extension of M1 where the genomic information is introduced via a covariance structure. Here, g_*i*_ represents the genomic effect of the *i*^*th*^genotype and it is computed as the linear combination between *p* markers (*X*_*im*_) and their respective marker effects (*b*_*m*_) {*m* = 1,2,…,*p*} such that 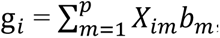, where 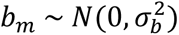 with 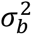 representing the corresponding variance component. Stacking all genomic into a single vector g = {g_*i*_} and by properties of the multivariate normal distribution we have that 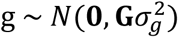 where 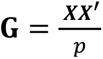 is the genomic relationship matrix describing genomic similarities between pairs of genotypes, ***X*** is the (standardized by columns) matrix of marker SNPs, and 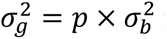 is the associated variance component (VanRaden, 2008). The resulting model can be represented as follows

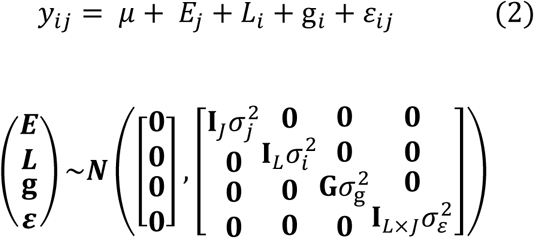

This model can be used to predict the non-phenotyped individuals as it allows the borrowing of information between individuals through the genomic relationship matrix.

### M3: E+L+G+GE [environment + genotype + genomic + genomic markers-by-environment (G×E) interaction]

In this model, the M2 is extended by adding the random term g*E*_*ij*_ which corresponds to the interaction between the *i*^*th*^ genotype and *j*^*th*^ environment. Stacking all the interaction effects into a single vector and following Jarquin et al., (2014) we have that 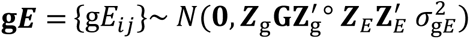 where 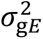 is the corresponding variance component, ***Z***_g_ and ***Z***_*E*_ are the design matrices connecting phenotypes to genotypes and environments, respectively, and “°” is the Hadamard product indicating cell-by-cell multiplication between two matrices of the same order. Thus, the model can be written as follows

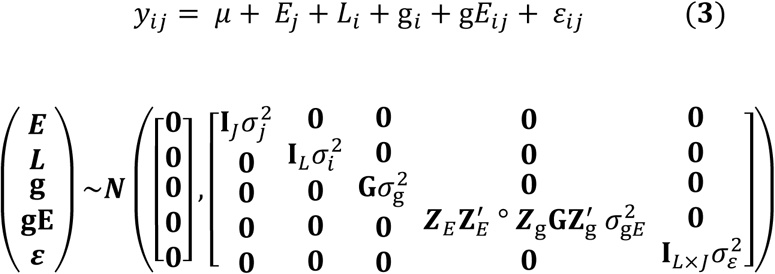

The previously mentioned model M2 returns common genetic effects for the same individuals across environments while with M3 the effects are particular to each environment.

### Cross-validations (CV)

To evaluate the prediction ability (PA) of the different sparse testing designs (Table 1) where the training sets varied their size and composition, the two principal cross-validation schemes (CV1 and CV2) were used as reference. The phenotypic data for the genotypes across environments was available, however, a fraction of the data was masked intentionally as missing values (NA) following different designs. As explained earlier, non-overlapping genotypes follow the basis of the CV2 scheme while overlapping genotypes follow the basis of the CV1 scheme. To elaborate, in the CV2 scheme, PA is driven by utilizing genomic similarities among genotypes and correlations between environments, thus borrowing phenotypic information from genotypes across environments. CV1 presents the situation where some genotypes are never tested in any of the environments and instead, predicting their performance relies genomic information and on other genotypes that were tested in fields.

The allocation designs (combinations of NO/O genotypes) in Table 1 show the gradual transitions between CV2 and CV1 modifying the numbers of NO and O genotypes in environments through random cross-validations. For example, the design 112/0 in Table 1 is a pure CV2 scheme while 0/112 is a pure CV1 scheme. In addition, with more NO and fewer O genotypes, the design closely aligns with the CV2 scheme. As the number of O genotypes increases and NO genotypes decrease across designs (from the left-hand side toward the right-hand side) the design gets closer to the CV1 scheme. Hence, Table 1 exhibits situations of both CV1 and CV2 schemes based on the numbers of NO/O genotypes across designs.

Here, the effects of the training compositions set sizes of the different sparse testing designs were assessed using the three above-mentioned prediction models. In all cases, PA was evaluated as the within-environment Pearson correlation between the predicted and the observed values of the unobserved 224 genotypes. In addition, MSE was also calculated on a trial basis where lower MSE values reflected a better performance of the models.

### Software

All the models were fitted using the BGLR package (Pérez and de los Campos, 2014) in R statistical programming language (R Core Team 2023).

## Results

The average results for PA and MSE across the three environments, 10 replicates of the different allocation designs, three prediction models, and four traits are presented in Figures 1 and 2. For each trait, the PA (Appendix Figures S2-S5) and MSE (Appendix Figures S6-S9) values were computed between observed and predicted values within each environment. Afterward, the mean average PA and MSE were calculated across environments by taking the mean of the within-environment values.

**Figure 1.**
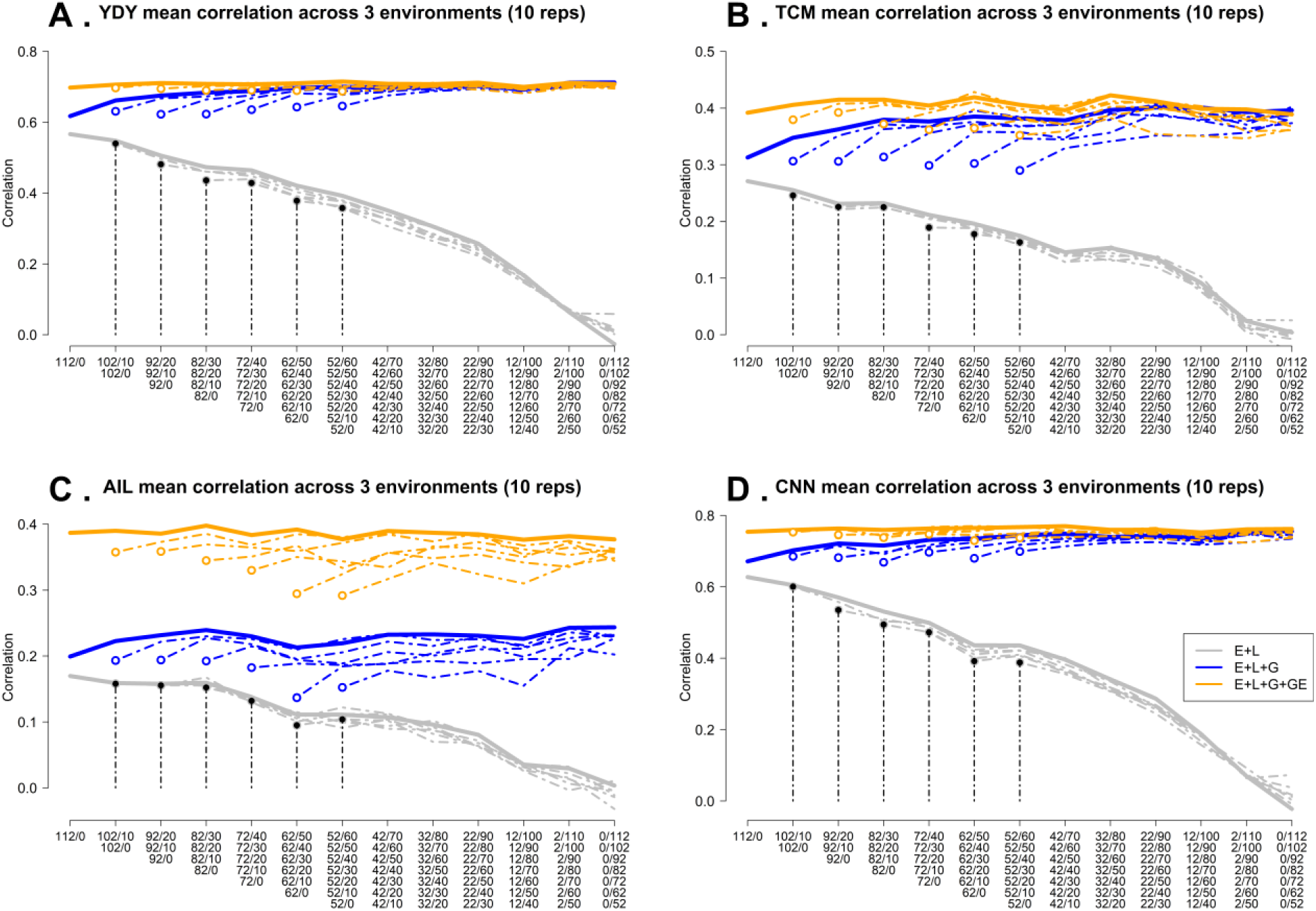
The mean PA (correlation between predicted and observed values) for four traits in varying training sets and allocation designs. A. dry biomass yield (YDY), B. total culm numbers (TCM), C. average internode lengths (AIL), D. culm node numbers(CNN) across three environments in three genomic prediction models, (M1**: E+L**; M2**: E+L+G**; M3**: E+L+G+GE**). Bold lines show the average PA for the largest calibration set (112 genotypes). Dotted lines show the reduced training sets (102, 92, 82, 72, 62, and 52 genotypes). X-axis shows the compositions of different non-overlapping (NO) / overlapping (O) allocation designs. Leftmost side of the X-axis shows compositions for completely NO genotypes with 0 overlaps across environments for different testing set sizes (e.g., 112/0 and 52/0). From left to right, the NO/O ratio decreases with increasing overlapping genotypes. Vertical, black-dashed lines indicate PA for different reduced training sets (e.g., 112/0, 92/0, and 52/0).

**Figure 2.**
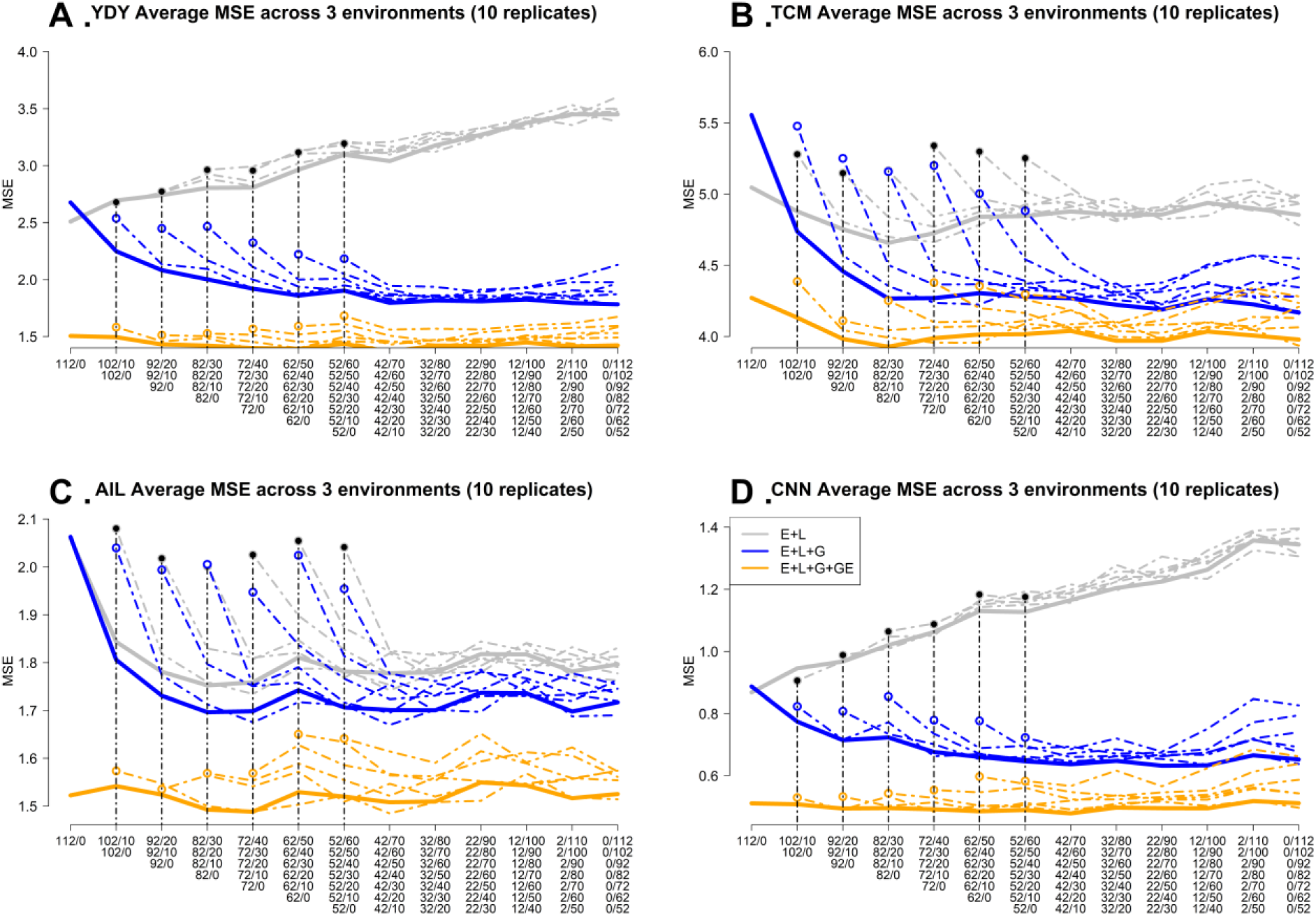
The mean MSE values for four traits in varying training sets and allocation designs. A. dry biomass yield (YDY), B. total culm numbers (TCM), C. average internode lengths (AIL), D. culm node numbers (CNN) across three environments in three genomic prediction models, (M1**: E+L**; M2**: E+L+G**; M3**: E+L+G+GE**). Bold lines show the average PA for the largest calibration set (112 genotypes). Dotted lines show the reduced training sets (102, 92, 82, 72, 62, and 52 genotypes). X axis shows the compositions of different non-overlapping (NO) / overlapping (O) allocation designs. Leftmost side of the X axis shows compositions for completely NO genotypes with 0 overlaps across environments for different testing set sizes (e.g., 112/0 and 52/0). From left to right, the NO/O ratio decreases with increasing overlapping genotypes. Vertical, black-dashed lines indicate PA for different reduced training sets (e.g., 112/0, 92/0, and 52/0).

### Predictive ability

For YDY, models M3 **(E+L+G+GE)** and M2 **(E+L+G)** always outperformed M1 **(E+L)** with M3 obtaining the highest PA (∼ 0.7) at the 112/0 design (Figure 1A). This result was higher around 13% and 23% compared to M2 (∼0.62) and M1 (∼0.58). The superiority of M3 remained consistent across all designs (from the left end at 112/0 towards the right end at 0/112). In other words, M3 was not influenced by the increasing number of overlapping genotypes across designs (middle as well as right sides of the plots). The PA in M2 slowly increased as more overlapping genotypes were added. For example, at designs 102/10, 92/20, and 72/40, the PA in M2 was ∼ 0.65, ∼0.66, and ∼0.68, respectively, which then plateaued at ∼0.70 the 40/72 design. M1 started with a PA ∼0.58 at 112/0, however, as expected, it significantly dropped towards 0 as the number of overlapping genotypes increased.

Considering the reduced training sets, M2 and M3 were not significantly impacted by reducing the sample size of the training sets. As expected, the PA in M1 reduced as the training set sizes decreased. In this case, the results maintained a decreasing trend with the rising number of overlapping genotypes across designs as well.

Similarly to the previous trait, there was a superiority of M3 concerning the PA over M2 and M1 for TCM (Figure 1B). The PA in M1 significantly reduced with the training set sizes. For M3 and M2, the PA ranged between ∼0.35 - ∼0.41 and ∼0.28 - ∼0.39, respectively [range between the smallest training set and the largest training set] for varying calibration set sizes. Moreover, for M2, the PA gradually increased as the number of overlapping genotypes. For example, the PAs in M2 at designs 112/0, 82/30, 32/80, and 12/100 were ∼0.31, ∼0.37, ∼0.38, and ∼0.39, respectively, for the full training set size. In contrast, M1 noticeably dropped after more overlapping genotypes were added. The reduced training set sizes slightly influenced both M2 and M3 results. In nearly all situations with a few exceptions, M3 delivered the best results over M2 under the same combinations of training sets and designs.

For AIL, M3 clearly outperformed both M2 and M1 across designs (Figure 1C). As more common genotypes were added, the PA did not significantly change for M2 and M3, while it gradually dropped for M1. In this case, the reduced training set sizes impacted the results of both M2 and M3 models. However, M3 was always superior for the same compositions of training sets and allocation designs over M2. The PA between the smallest and the largest training set ranged from ∼0.28 to∼0.41 for M3 and from ∼0.13 to ∼0.23] for M2. As expected and similar than for the other traits, M1 was significantly impacted with reduced training set sizes, as well as for the increasing number of overlapping genotypes.

Finally, similar patterns regarding the performance of M1, M2, and M3 for all compositions of training sets and allocation designs were observed for CNN (Figure 1D). Both M2 and M3 performed better compared to M1, however, M3 achieved the highest PA ∼0.77 at 112/0 and remained unchanged as the number of NO/O ratios reduced (as the number of overlapping genotypes increased). For M2, the PA gradually increased with decreasing NO/O. For example, M2 obtained a PA of ∼0.67 at 112/0, ∼0.70 at 72/40, and by 2/102, and it reached a maximum plateau of ∼ 0.77. However, although the best results of M2 are comparable to those from M3 those from the latter do not require a high number of overlapping genotypes. In contrast, M1 always returned the lowest PA across all designs, showing a gradual drop as the number of overlapping genotypes was increased.

In summary, consistently M3 returned the best results while as expected M1 showed the worst results for all traits in all compositions of calibration set sizes and allocation designs. For YDY and CNN, M2 and M3 showed smaller differences between these than for the other traits with M3 outperforming M2 considering the larger number of non-overlapping genotypes in the designs. In addition, for all traits the PA remained almost unchanged with M3 as more genotypes were added to the designs. For M2, the PA slowly increased as the number of overlapping genotypes increased as well. While for M1 it was consistently reduced as the ratio of NO/O decreased. In general, accuracy was lower in AIL and TCM compared to YDY and CNN. For AIL, the PA obtained with M3 and M2 was negatively affected by reducing the training set sizes but remained stable for YDY, TCM and CNN.

### Mean square error

Figures 2A, 2B, 2C, and 2D display the average MSE for YDY, TCM, AIL, and CNN respectively for M1 **(E+L)**, M2 **(E+L+G)**, and M3 **(E+L+G+GE)** models across all environments and 10 replicates. Here, the performance of the models with the smallest MSE is considered the best. For YDY, both M3 (∼1.5) and M2 (∼2.4) performed better than M1 (2.5) with M3 consistently returning the lowest MSE values (Figure 2A). Moreover, M3 remained constant as the ratio of NO/O reduced across designs. At the beginning (112/0) when the genotypes were tested only once, the MSE value in M1 was slightly lower (∼2.5) compared to M2 (∼2.7) (Figure 2A). However, as more common genotypes were added to the designs, the values gradually increased for M1 while decreased for M2. With the reduced calibration set sizes, the MSE values in M3 increased slightly to ∼1.7, while M1 significantly increased. Interestingly, with M2 the MSE values slightly reduced. Nevertheless, M3 always returned the lowest values compared to M2 and M1 across all NO/O genotype compositions (from the left-hand side towards the right-hand side).

Similar to the previous trait, M3 achieved the smallest MSE value for TCM. The values slightly varied between ∼3.8 and ∼4.3 across designs (Figure 2B). M1 had a slightly lower value (∼5.1) at 112/0 compared to M2 (∼5.6). However, for different compositions of NO/O genotypes, the MSE reduced in both M2 (∼4.8) and M1 (∼4.3) with M2 experiencing a more pronounced reduction. When the calibration set sizes reduced, all models were impacted where M1 and M2 showed a decreasing pattern. On the other hand, despite M3 showing a slightly increasing pattern in MSE, it constantly yielded the best results with lower MSE than M1 and M2.

For AIL, the MSE was always low with M3 compared to M1 and M2 (Figure 2C). The values for M3 ranged between ∼1.48 to ∼1.52 for different combinations of NO/O genotypes. For models M1 and M2 both started with a MSE value of ∼2.08 at 112/0; however, for M2 it declined for the subsequent allocation schemes (designs in the middle and on the right-hand side of the plots) as the number of common genotypes increased. Although both M1 and M2 decreased, the reduction was more pronounced in M2 compared to M1. With reduced training set sizes, all models were impacted for different compositions of NO/O. The MSE values slightly increased in all three models with M1 and M2 showing a decreasing pattern while M3 showing a slightly increasing pattern. Despite the increasing trajectory observed in M3 it always returned the best results.

Finally for CNN, the models demonstrated similar trends as observed in YDY (Figure 2D). M3 constantly produced the lowest MSE (∼0.5) compared to both M2 and M1. The prominence of M3 was shown for all combinations of NO/O ratio. On the other hand, M1 obtained a MSE value of ∼0.88 while M2 of ∼0.9 at the design 112/0. With the reduced NO/O ratio, the value kept increasing in M1, whereas decreasing in M2. Reduced training set sizes resulted in a marginal increase of the MSE value for M3 (∼0.62). However, despite the minimal change observed in M3, it always outpaced both M1 and M2 across the comparable combinations of training sets and designs.

Although, in general, M2 and M3 both performed better than M1 in all traits, M3 always achieved the lowest MSE values for all compositions of calibration set sizes and allocation designs. Also, the MSE values varied minimally in M3 when more overlapping genotypes were added across designs.

Overall, the results in PA and MSE suggest that M3 was the most superior model in all traits for different compositions of calibration sets and sparse testing designs. In addition, M3 remained constant as the number of overlapping genotypes increased across designs. Although in some cases, the results in M3 were slightly influenced by the reduced sample sizes based on traits, it still outperformed both M2 and M1. On the other hand, M1 returned the worst results; while, for M2 the results improved with the addition of common genotypes.

Therefore, the results indicate that evaluating different fractions of non-overlapping *Miscanthus* genotypes in all environments or repeating some overlapping genotypes in all environments would generate almost similar accuracy. The performance of M3 shows that either approach would be effective. However, the accuracy would also depend on the genetic architecture of the traits including the types of environments they are evaluated. The observed results show that there is no optimum fraction for using the number of overlapping genotypes. However, a small set of 10 to 20 overlapping genotypes could be better for research purposes to estimate environmental variability that is not confounded with genetic variability (Jarquin et al., 2020).

## Discussion

The development of improved varieties requires establishing multi-environment trials (METs) to evaluate their performance under a wide range of environmental conditions (Brown et al., 2020; Smith et al., 2021). METs play a crucial role in effectively capturing G×E interaction by evaluating multiple genotypes in different environments (Verbyla, 2023; Argaw et al., 2025). However, setting up METs in *Miscanthus* is resource-intensive as the establishment cost due to rhizomatous breeding is high (Arnoult and Brancourt-Hulmel, 2015). Additionally, prolonged field testing that requires *Miscanthus* crops to wait for three years to yield full potential is required for phenotyping (O’Loughlin et al., 2017), which lengthens the release of commercial *Miscanthus* varieties. Hence, it is difficult to determine the strategic selection and allocation of superior *Miscanthus* genotypes across target environments.

Genomic selection has emerged as a powerful tool to inform breeders about the selection of genotypes in a more effective way, with the initial goal of boosting genetic gain (Budhlakoti et al., 2022; Alemu et al., 2024). In addition, it plays a vital role in addressing challenges with complex traits and G×E confronted in METs. Previous studies have explored different ways of incorporating G×E into the GP models (Jarquin et al., 2014; Acosta-Pech et al., 2017). González-Barrios et al., (2019) explored resource optimization across multiple environments by using mega-environments while including G×E into the model. In the *Miscanthus* population, only a few studies have leveraged the use of GP models in METs (Slavov et al., 2014; Clark et al., 2019; Dong et al., 2019; Njuguna et al., 2023).

Clark et al., (2012) and Olatoye et al., (2020) showed the importance of relatedness between *Miscanthus* individuals while designing a training population for use in GP models. However, no previous studies have explored the effects of sparse testing allocation strategies on the accuracy of GP models for *Miscanthus* METs using different combinations of non-overlapping and overlapping genotypes. As with other perennial crops, GP-based sparse testing is of particular interest in *Miscanthus* METs because initial evaluation of a large number of *Miscanthus* genotypes requires high establishment and phenotyping costs across multiple years (Lewandowski et al., 2016). Optimum allocation of the superior *Miscanthus* genotypes can reduce the total costs of the initial trials.

In this study that considers 336 genotypes, three environments, and 1,008 phenotypic records, it was shown that GP-based sparse testing strategies could maintain high accuracy at different compositions of calibration sets. Hence, a substantial reduction of phenotyping costs can be accomplished, especially when using model M3. In addition, the accuracy remained practically unchanged as more overlapping genotypes were added to the same model. Thus, the incorporation of G×E into prediction models has the potential to take information from genotypes observed in target environments and apply it to novel environments. However, maintaining consistent PA while adding more overlapping genotypes has less significance in *Miscanthus* breeding in North America, considering costs. This is because *Miscanthus* relies on vegetative propagation where the cost is initially associated with establishments, demanding intensive labor, management, and maintenance. Therefore, in *Miscanthus* breeding, the focus on cost-saving mainly shifts toward observing fewer genotypes while maintaining high PA in the target environments by optimizing resource allocation.

The results regarding the decreasing calibration set sizes showed that PA remained almost unchanged across all traits, especially while using model M3. This model provided an opportunity to increase the testing capacity of evaluating more genotypes by ensuring similar accuracy, thus saving costs associated with resource management. For example, evaluating 52 genotypes in the smallest training set in each of the three environments resulted in almost the same accuracy level as with 112 genotypes in the largest training set (336/3=112), while using model M3. Thus, instead of observing a total of 1008 phenotypes for 336 genotypes-in-environment combinations (336×3=1008), evaluating only 52 genotypes (52×3=156 phenotypes) in each of the three environments would reduce the phenotyping costs by ∼85% [= (1008-156)/1008]. Obviously, the actual cost in real *Miscanthus* breeding programs would vary from this estimate since also differences in cost, labor, management, etc. vary among environments/locations. However, evaluating fewer genotypes would still save the costs associated with phenotyping data collection, and environmental inputs such as water use, fertilizer use, etc.

Albeit overlapping genotypes have minor importance in *Miscanthus* breeding, adding a small fraction could be useful for research intent. It could act as a baseline for comparison while estimating environmental variance across different environments. This is because, in the models, the overlapping genotypes behave as a control while separating environmental variance from the genetic variance. Another reason for adding some overlapping genotypes could be logistical restraints that limit testing genotypes in some environments but allow evaluation of them in others (Jarquin et al., 2020). Hence, including a fraction of 10 to 20 overlapping genotypes with the rest of the non-overlapping genotypes (42) distributed in three environments would be an effective approach for substantial cost savings in *Miscanthus* breeding.

## Comparison of the prediction models

An important objective of this study was to investigate the effects of three GP models for varying training set sizes and sparse testing designs across three environments and four traits. The model that returned maximum PA and minimum MSE for different combinations of NO/O genotypes for varting training sets was considered the best. Results indicated that model M1 had the lowest PA and highest MSE compared to M2 and M3. When the genotypes were observed in at least one environment, the PA was better in M1 (112/0 or 52/0). This is because, when a genotype is observed in at least one environment, the effect of genotypes is blended with the effect of environments.

The phenotypic information of the observed genotypes had therefore a strong impact on GP accuracy. However, the accuracy rapidly dropped across designs when the ratio of NO/O reduced (from the left extremes to the right extremes). It is because when the ratio of NO/O reduced, the design slowly approached towards the true CV1 scheme by 0/112. In CV1, there is no observed information available on the genotypes, and it is impossible for M1 to borrow information for the observed genotypes. As no marker information was included in M1, it could not borrow information on the genotypes through genomic similarities. Similar patterns were found in Jarquin et al. (2020) and Garcia-Abadillo et al. (2024) for maize and sugarcane datasets respectively where prediction accuracy decreased for M1 in the situation where no phenotypic information was available for the genotypes. On the other hand, MSE in M1 highlighted the opposite pattern, increasing with reduced NO/O fractions.

For all traits, M3 had the highest PA. The main reason is that M3 included G×E interaction in the model which could borrow information about the behavior of the genetic value of the genotypes across and within environments. Although M2 and M3 had nearly similar accuracies in YDY and CNN, the MSE value was lower in M3 for the same compositions of training sets and NO/O combinations. This may be attributable to the G×E term used in M3, which could significantly reduce the unexplained proportion of the total variance compared to M2. Hence, constant low MSE values in M3 compared to M2 suggested that it was the best prediction model for *Miscanthus* sparse testing designs.

Another notable trend in M3 was that it consistently returned almost the same accuracy across designs in all traits. In other words, adding more overlapping genotypes did not change the accuracy in M3. To elaborate, with the addition of overlapping genotypes, the designs shift towards CV1 where there is a lack of phenotyping information on the untested genotypes, hence depending on genomic marker data to fill missing information. In this circumstance, the G term in M3 model could borrow information of the genotypes via genomic similarities whereas the G×E term could capture the interaction more effectively. This finding also suggests that for the *Miscanthus* population, it does not matter whether complete non-overlapping genotypes are tested in each environment or if some overlapping sets are used across environments. In both cases, it is possible to reduce the phenotyping cost; however, the accuracy could vary depending on the traits and how correlated the environments are between them. Also, as discussed although for same TRAINING set size similar results were obtained with the varying compositions, some of these are more plausible to occur in the real application in breeding programs where availability of materials could be a bottleneck for propagation.

These findings regarding constant PA across designs in *Miscanthus* differ from the observations reported in sugarcane (Garcia-Abadillo et al., 2024). In sugarcane, the PA decreased in the M3 model as more overlapping genotypes were added. This could be due to a strong environmental correlation or due to including small sample sizes in the training sets. Considering the PA across traits, it was observed that M3 returned higher PA in YDY and CNN compared to AIL and TCM. Moreover, decreasing the calibration set size did not influence the accuracies in YDY and CNN but negatively impacted TCM and AIL. Also, the MSE varied minimally for YDY and CNN but varied significantly for TCM and AIL with reduced training set size in M3. With full calibration set size, the PA in TCM and AIL was ∼0.41. This value was reduced by ∼18 to ∼ 42% respectively in TCM and AIL as the training set was reduced. Njuguna et al., (2023) reported prediction accuracies in the same *Miscanthus* population for YDY, CNN, AIL, and TCM that were 0.56, 0.53, 0.15, and 0.46, respectively. The key difference between this study and our study is that we used sparse testing designs with 33% of the genotypes in the TRAINING sets and 66% of the genotypes for the testing set.

In contrast, Njuguna et al., (2023) used more phenotypic records for their analysis. The author also stated the importance of a large training population for high prediction accuracies in the *M. sacchariflorus* population. Hence, TCM and AIL could require a large training set size (> 112) to capture the variability for improved prediction accuracy. In contrast, for YDY and CNN, the G×E interaction term could account for most of the variances in a smaller training set size (52) and return similar accuracies as with the largest training set size (112). Another key aspect of the results obtained in this study was the use of high-density markers. For the present study, a total of around 136k was available for performing GP analysis. In practical situations, it is important to account for budget constraints that could be associated with genotyping costs along with allocation schemes. Jarquin et al., (2020) used ∼62k makers for deploying sparse testing designs in the maize dataset and found reasonable accuracies yielded by M3.

In sugarcane, which is a closest species of *Miscanthus*, ∼22k markers were enough for M3 to return PA varying between ∼0.34-0.61 (Garcia-Abadillo et al., 2024). However, reducing marker density could lower accuracy for conducting sparse testing designs in *Miscanthus*. Thus along with allocation strategies, it is important to balance between marker density and PA to maximize genetic gain and cost-efficiency. If accuracy is reduced with fewer markers, alternative approaches will need to be considered in the genomic prediction models.

## Practical implementation of sparse testing in *Miscanthus* breeding

*Miscanthus* has a broad geographical distribution related to a diverse set of germplasms with multiple adaptation ranges across Asia to Siberia (Huang et al., 2019). It is an excellent source for establishing *Miscanthus* breeding programs in North America. However, there are various challenges to identifying and selecting the best variety for the breeding programs (Clifton-Brown et al., 2019). Besides finding the best crosses, evaluating *Miscanthus* genotypes requires expensive rhizomatous or in vitro propagation for the establishment (Xue et al., 2015). Given the breeding trials, previous studies in *Miscanthus* focused mainly on the yield and yield component traits (Kaiser et al., 2015; Clark et al., 2019; Njuguna et al., 2023). However, extensive multilocation field trials are expensive as most yield and yield component traits cannot be measured before two to three years of establishment. There are additional costs for labor, land (Lewandowski et al., 2016), and genotyping for optimizing molecular marker use. Therefore, finding the superior genotypes and reducing the associated costs are important issues to consider while increasing genetic gain in *Miscanthus* breeding programs.

Considering *Miscanthus* breeding programs with fixed budgets and the long establishment phages, the breeders must carefully decide at the initial stages how many genotypes they should evaluate in fields. On the other hand, they should also optimize genotyping procedures for the rest of the genotypes and aim for efficient resource allocations to maximize genetic gain. The practical implementation of GP-based sparse testing in *Miscanthus* breeding shows how to optimize resource allocation while maintaining high PA and decreasing phenotyping costs by adjusting varying compositions of NO/O genotypes. With the saved costs, more elite genotypes can be evaluated, thus increasing the selection intensity and eventually optimizing genetic gain.

This study in sparse testing also highlights how enhancing selection intensity by increasing the evaluation of genotypes eventually leads to improved genetic gain. Although this study did not assess how an increase in selection intensity would maximize genetic gain, it still showed the potential of evaluating more genotypes by allocating resources as a factor in optimizing genetic gain. Overall, this study aligns with this goal by illustrating how G×E components in the GP model behave under different compositions of non-overlapping and overlapping genotypes. By finding compositions that maximize accuracy, breeders can better target superior *Miscanthus* genotypes with well-adaptability, increasing final genetic gains.

## Conclusions

Genome-based sparse testing design is an approach to accelerate breeding programs cost-effectively. It allows testing candidate genotypes in many environments and provides a detailed evaluation regarding their broad adaptability and stability. In the present study, different compositions of training sets and allocation schemes were assessed in *M. sacchariflorus* multi-environment trials. The GP M3 **(E+L+G+GE)** model that incorporated the G×E interaction term, showed the best PA with low MSE across allocation designs for all the traits. It denotes that incorporating G×E in GP model improved the PA and can be utilized in sparse testing.

Overall, M3 provided an opportunity to maintain high accuracy even after reducing the calibration set sizes. This could significantly reduce phenotyping costs in multi-environment *Miscanthus* breeding trials. Also, it can increase the testing capacity of the *Miscanthus* genotypes in METs. From this study, an ideal design strategy for multi-environment *Miscanthus* trials could involve testing a few common genotypes along with more non-overlapping genotypes assigned to all environments.

## Acknowledgements

This work was funded by the DOE Center for Advanced Bioenergy and Bioproducts Innovation (U.S. Department of Energy, Office of Science, Office of Biological and Environmental Research under Award Number DE-SC0018420). Any opinions, findings, and conclusions or recommendations expressed in this publication are those of the authors and do not necessarily reflect the views of the U.S. Department of Energy.

## Conflict of interest

The authors declare no conflict of interest

## Data Availability

The data sets and analyses implemented in this research can be found in https://doi.org/10.6084/m9.figshare.31796794

## Appendix

### Figures S1

**Figure S1.**
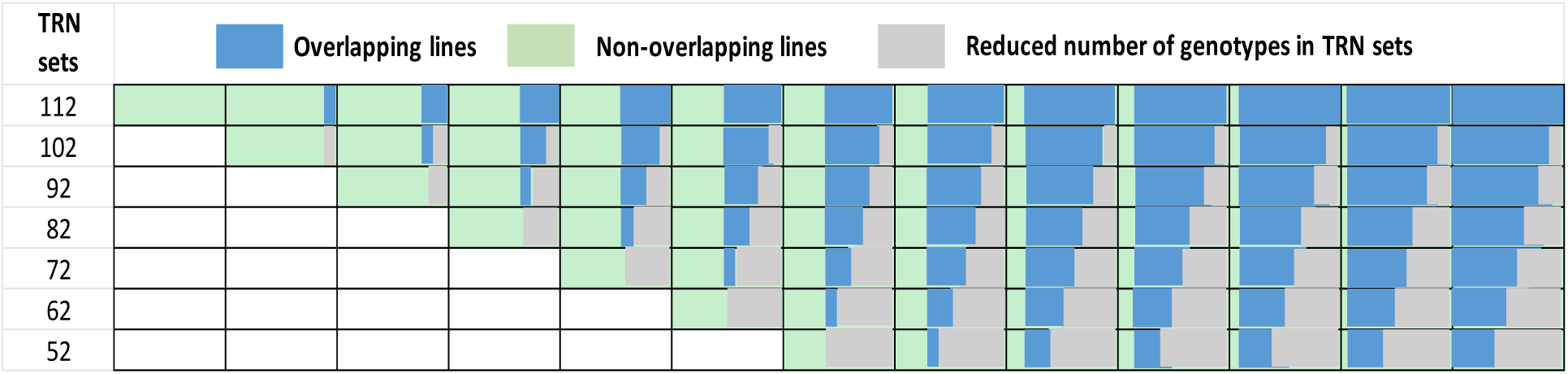
Graphical presentation of different allocation schemes. The rows present different TRAINING sets (from 112 until 52) and the columns present the allocation designs. The blue color shows the overlapped sets, light green presents the non-overlapping sets and the grey color displays the reduced TRAINING sets. Across designs (from left to right), the number of non-overlapping genotypes decreases while the number of overlapping genotypes increases.

### Figures S2-S5

The PA between predicted and observed value for each trait within each environment.

**Figure S2.**
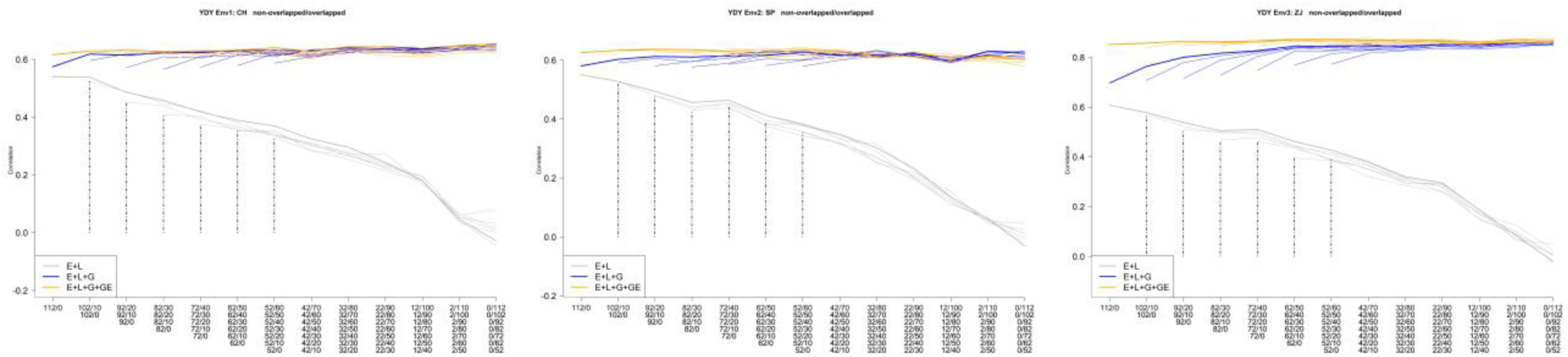
The correlation between predicted and observed values for varying TRAINING sets and allocation designs in YDY [Left: correlation for YDY in environment 1, Middle: correlation for YDY in environment 2, Right: correlation for YDY in environment 3] within each environment in three genomic prediction models, M1: **E+L;** M2: **E+L+G;** M3: **E+L+G+GE**. Bold lines show the average PA for the largest calibration set (112 genotypes). Dotted lines show the reduced training sets (102, 92, 82, 72, 62, and 52 genotypes). X axis shows the compositions of different non-overlapping (NO) / overlapping (O) allocation designs. Leftmost side of the X axis shows compositions for completely NO genotypes with 0 overlaps across environments for different testing set sizes (e.g., 112/0 and 52/0). From left to right, the NO/O ratio decreases with increasing overlapping genotypes. Vertical, black-dashed lines indicate PA for different reduced training sets (e.g., 112/0, 92/0, and 52/0).

**Figure S3.**
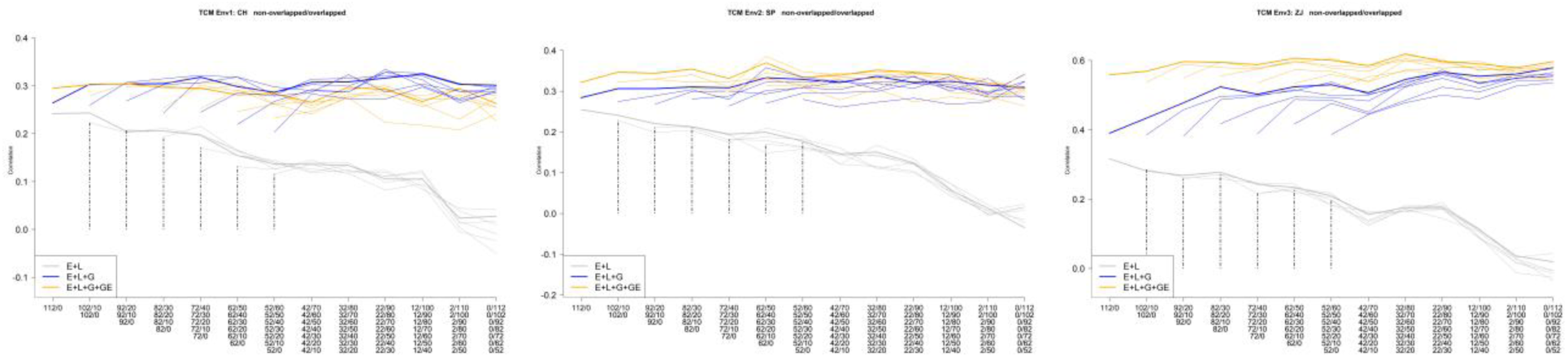
The correlation between predicted and observed values for varying TRAINING sets and allocation designs in TCM [Left: correlation for TCM in environment 1, Middle: correlation for TCM in environment 2, Right: correlation for TCM in environment 3] within each environment in three genomic prediction models, M1: **E+L;** M2: **E+L+G;** M3: **E+L+G+GE**. Bold lines show the average PA for the largest calibration set (112 genotypes). Dotted lines show the reduced training sets (102, 92, 82, 72, 62, and 52 genotypes). X axis shows the compositions of different non-overlapping (NO) / overlapping (O) allocation designs. Leftmost side of the X axis shows compositions for completely NO genotypes with 0 overlaps across environments for different testing set sizes (e.g., 112/0 and 52/0). From left to right, the NO/O ratio decreases with increasing overlapping genotypes. Vertical, black-dashed lines indicate PA for different reduced training sets (e.g., 112/0, 92/0, and 52/0).

**Figure S4.**
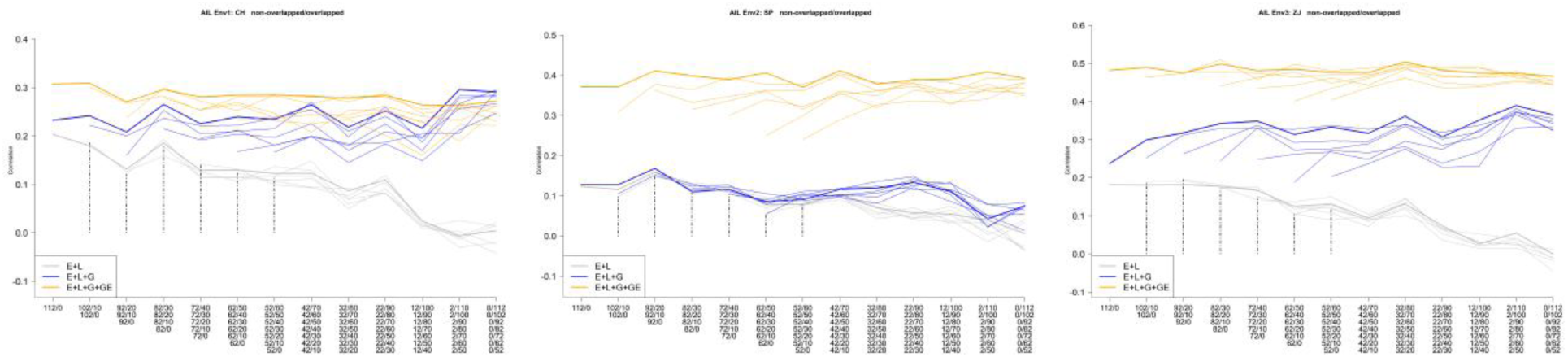
The correlation between predicted and observed values for varying TRAINING sets and allocation designs in AIL [Left: correlation for AIL in environment 1, Middle: correlation for AIL in environment 2, Right: correlation for AIL in environment 3] within each environment in three genomic prediction models, M1: **E+L;** M2: **E+L+G;** M3: **E+L+G+GE**. Bold lines show the average PA for the largest calibration set (112 genotypes). Dotted lines show the reduced training sets (102, 92, 82, 72, 62, and 52 genotypes). X axis shows the compositions of different non-overlapping (NO) / overlapping (O) allocation designs. Leftmost side of the X axis shows compositions for completely NO genotypes with 0 overlaps across environments for different testing set sizes (e.g., 112/0 and 52/0). From left to right, the NO/O ratio decreases with increasing overlapping genotypes. Vertical, black-dashed lines indicate PA for different reduced training sets (e.g., 112/0, 92/0, and 52/0).

**Figure S5.**
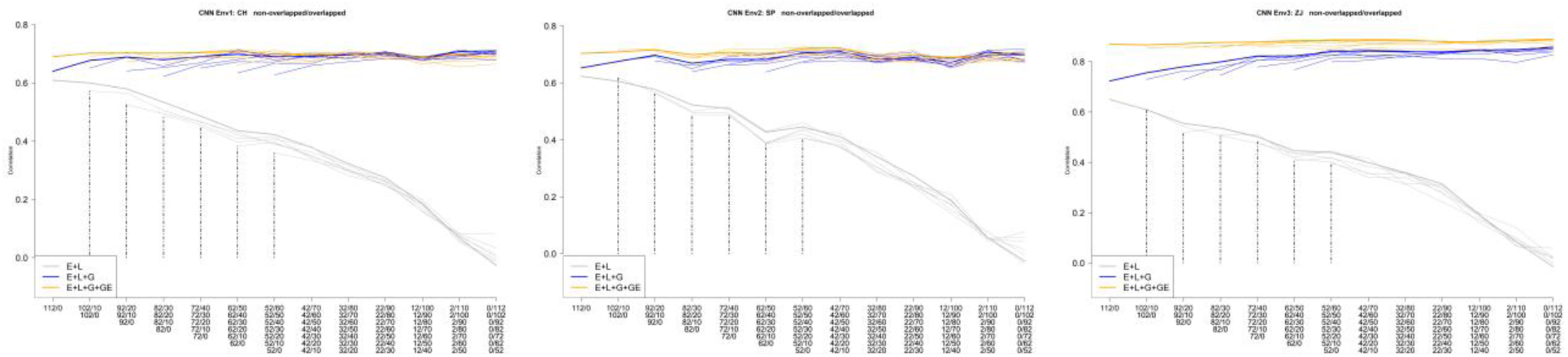
The correlation between predicted and observed values for varying TRAINING sets and allocation designs in CNN [Left: correlation for CNN in environment 1, Middle: correlation for CNN in environment 2, Right: correlation for CNN in environment 3] within each environment in three genomic prediction models, M1: **E+L;** M2: **E+L+G;** M3: **E+L+G+GE**. Bold lines show the average PA for the largest calibration set (112 genotypes). Dotted lines show the reduced training sets (102, 92, 82, 72, 62, and 52 genotypes). X axis shows the compositions of different non-overlapping (NO) / overlapping (O) allocation designs. Leftmost side of the X axis shows compositions for completely NO genotypes with 0 overlaps across environments for different testing set sizes (e.g., 112/0 and 52/0). From left to right, the NO/O ratio decreases with increasing overlapping genotypes. Vertical, black-dashed lines indicate PA for different reduced training sets (e.g., 112/0, 92/0, and 52/0).

### Figures S6-S9

The MSE values for each trait within each environment.

**Figure S6.**
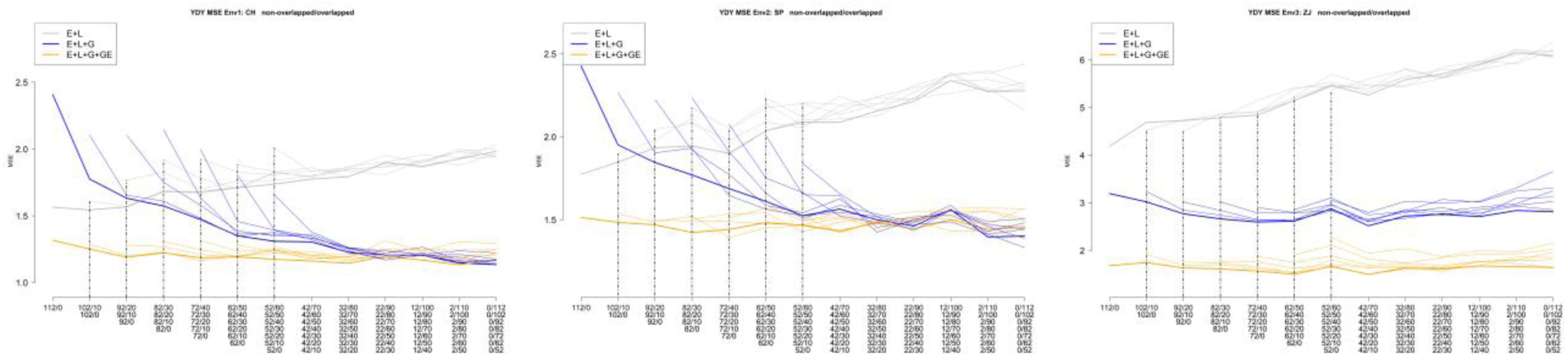
The MSE values for varying TRAINING sets and allocation designs in YDY [Left: MSE for YDY in environment 1, Middle: MSE for YDY in environment 2, Right: MSE for YDY in environment 3] within each environment in three genomic prediction models, M1: **E+L;** M2: **E+L+G;** M3: **E+L+G+GE**. Bold lines show the average PA for the largest calibration set (112 genotypes). Dotted lines show the reduced training sets (102, 92, 82, 72, 62, and 52 genotypes). X axis shows the compositions of different non-overlapping (NO) / overlapping (O) allocation designs. Leftmost side of the X axis shows compositions for completely NO genotypes with 0 overlaps across environments for different testing set sizes (e.g., 112/0 and 52/0). From left to right, the NO/O ratio decreases with increasing overlapping genotypes. Vertical, black-dashed lines indicate PA for different reduced training sets (e.g., 112/0, 92/0, and 52/0).

**Figure S7.**
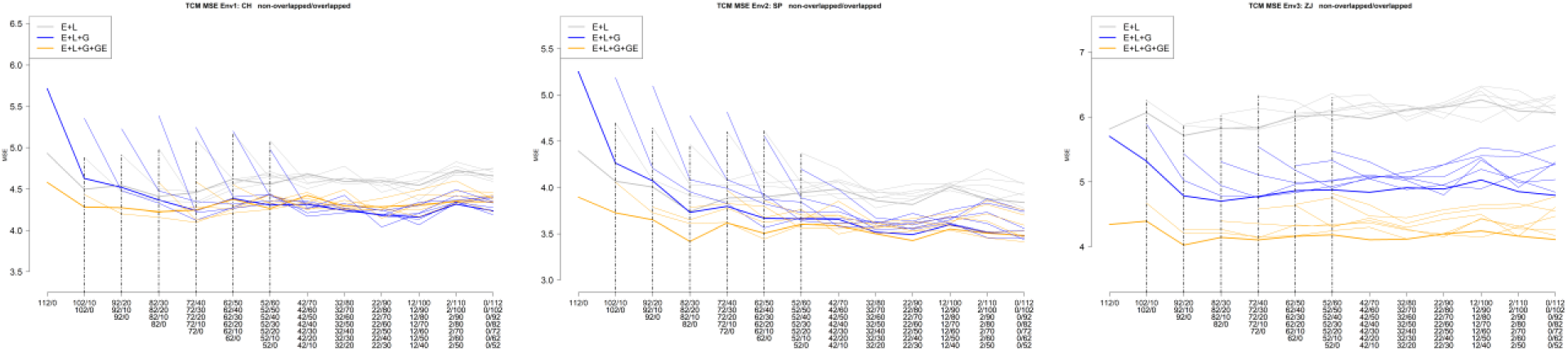
The MSE values for varying TRAINING sets and allocation designs in TCM [Left: MSE for TCM in environment 1, Middle: MSE for TCM in environment 2, Right: MSE for TCM in environment 3] within each environment in three genomic prediction models, M1: **E+L;** M2: **E+L+G;** M3: **E+L+G+GE**. Bold lines show the average PA for the largest calibration set (112 genotypes). Dotted lines show the reduced training sets (102, 92, 82, 72, 62, and 52 genotypes). X axis shows the compositions of different non-overlapping (NO) / overlapping (O) allocation designs. Leftmost side of the X axis shows compositions for completely NO genotypes with 0 overlaps across environments for different testing set sizes (e.g., 112/0 and 52/0). From left to right, the NO/O ratio decreases with increasing overlapping genotypes. Vertical, black-dashed lines indicate PA for different reduced training sets (e.g., 112/0, 92/0, and 52/0).

**Figure S8.**
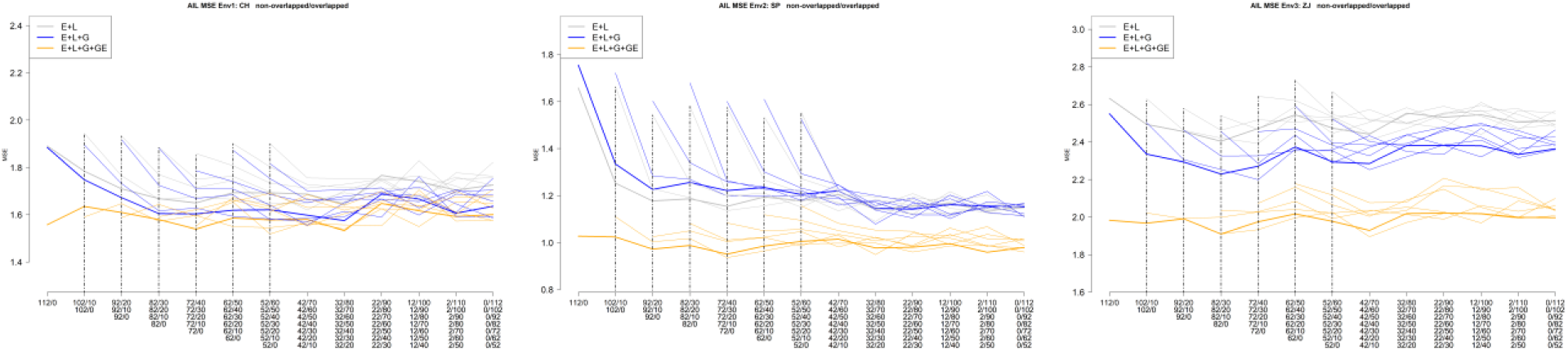
The MSE values for varying TRAINING sets and allocation designs in AIL [Left: MSE for AIL in environment 1, Middle: MSE for AIL in environment 2, Right: MSE for AIL in environment 3] within each environment in three genomic prediction models, M1: **E+L;** M2: **E+L+G;** M3: **E+L+G+GE**. Bold lines show the average PA for the largest calibration set (112 genotypes). Dotted lines show the reduced training sets (102, 92, 82, 72, 62, and 52 genotypes). X axis shows the compositions of different non-overlapping (NO) / overlapping (O) allocation designs. Leftmost side of the X axis shows compositions for completely NO genotypes with 0 overlaps across environments for different testing set sizes (e.g., 112/0 and 52/0). From left to right, the NO/O ratio decreases with increasing overlapping genotypes. Vertical, black-dashed lines indicate PA for different reduced training sets (e.g., 112/0, 92/0, and 52/0).

**Figure S9.**
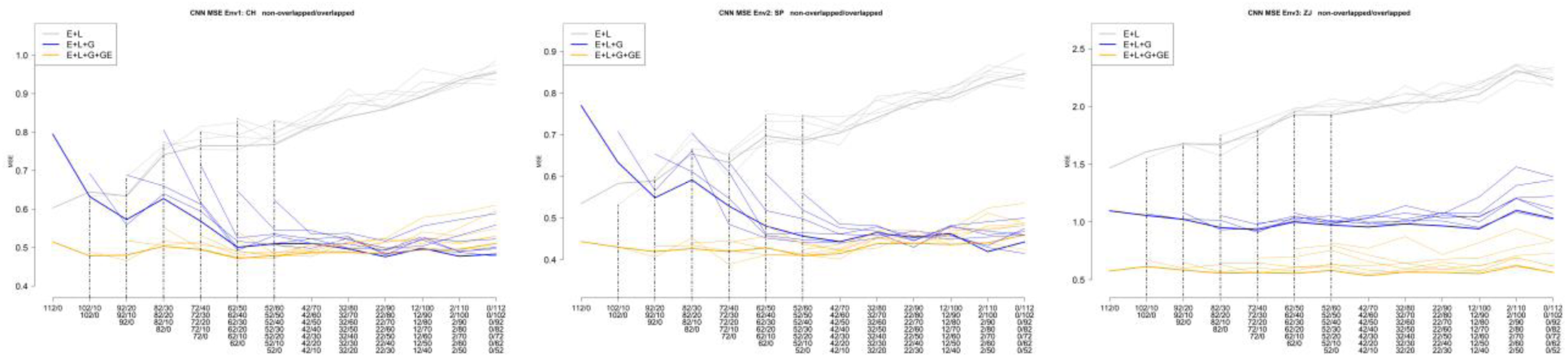
The MSE values for varying TRAINING sets and allocation designs in CNN [Left: MSE for CNN in environment 1, Middle: MSE for CNN in environment 2, Right: MSE for CNN in environment 3] within each environment in three genomic prediction models, M1: **E+L;** M2: **E+L+G;** M3: **E+L+G+GE**. Bold lines show the average PA for the largest calibration set (112 genotypes). Dotted lines show the reduced training sets (102, 92, 82, 72, 62, and 52 genotypes). X axis shows the compositions of different non-overlapping (NO) / overlapping (O) allocation designs. Leftmost side of the X axis shows compositions for completely NO genotypes with 0 overlaps across environments for different testing set sizes (e.g., 112/0 and 52/0). From left to right, the NO/O ratio decreases with increasing overlapping genotypes. Vertical, black-dashed lines indicate PA for different reduced training sets (e.g., 112/0, 92/0, and 52/0).

## Notes

### Competing Interest Statement

The authors have declared no competing interest.

